# Contrasting the genetic architecture of 30 complex traits from summary association data

**DOI:** 10.1101/035907

**Authors:** Huwenbo Shi, Gleb Kichaev, Bogdan Pasaniuc

**Affiliations:** Bioinformatics Interdepartmental Program, University of California, Los Angeles, 90024; Dept of Pathology and Laboratory Medicine; Dept of Human Genetics, David Geffen School of Medicine, University of California, Los Angeles, 90024

## Abstract

Variance components methods that estimate the aggregate contribution of large sets of variants to the heritability of complex traits have yielded important insights into the disease architecture of common diseases. Here, we introduce new methods that estimate the total variance in trait explained by a single locus in the genome (local heritability) from summary GWAS data while accounting for linkage disequilibrium (LD) among variants. We apply our new estimator to ultra large-scale GWAS summary data of 30 common traits and diseases to gain insights into their local genetic architecture. First, we find that common SNPs have a high contribution to the heritability of all studied traits. Second, we identify traits for which the majority of the SNP heritability can be confined to a small percentage of the genome. Third, we identify GWAS risk loci where the entire locus explains significantly more variance in the trait than the GWAS reported variants. Finally, we identify 55 loci that explain a large proportion of heritability across multiple traits.

## Introduction

Large-scale genome-wide association studies (GWAS) have identified thousands of single nucleotide polymorphisms (SNPs) associated with hundreds of traits^1,2,3,4^. However, only a fraction of the variance in trait can be explained by the risk SNPs reported by GWAS. The so-called “missing heritability problem” is in part due to the stringent significance threshold imposed in GWAS, which neglects causal variants that fail to reach the genome-wide significance level at current sample sizes. As an alternative, variance component (heritability) analysis aggregates the effect of all SNPs regardless of their significance to increase accuracy^5^ and has yielded important insights into the genetic architecture of complex traits^6,7,8,9,10,11^.

Standard approaches for heritability estimation and partitioning rely on estimating the genetic relationships between pairs of individuals (estimated genome-wide or across a subset of the genome)^8,12,13^. Therefore, these analyses require individual-level genotype data which prohibits their applicability to ultra-large GWAS that, due to privacy concerns, is typically available only at the summary level. To solve this bottleneck, recent methods have shown that heritability, both genome-wide as well as for different functional categories in the genome, can be accurately estimated using only summary GWAS data^6,7^. Although these methods have enabled powerful analyses making insights into genetic basis of complex traits, they rely on the infinitesimal model assumption (i.e. all SNPs contribute to the trait) which is invalid at most risk loci^6,7^. To overcome this drawback, alternative approaches have proposed to impose a prior on the sparsity of effect sizes to further increase heritability estimation accuracy^14^. A potentially more robust approach is to not assume any distribution for the effect sizes at causal variants and treat them as fixed effects in the heritability estimation procedure. Indeed, recent works have shown that heritability estimation can be performed under maximum-likelihood from polygenic scores under a fixed-effect model assuming no LD among SNPs^11^.

Here, we introduce Heritability Estimator from Summary Statistics (HESS), an approach to estimate the variance in trait explained by a single locus while accounting for linkage disequilibrium (LD) among SNPs. We build upon recent works^11,15^ that treat causal effect sizes as fixed effects and model the genotypes at the locus as random correlated variables. Our estimator can be viewed as a weighted summation of the squares of the projection of GWAS effect sizes onto the eigenvectors of the LD matrix at the considered locus, where the weights are inversely proportional to the corresponding eigenvalues. Through extensive simulations, we show that HESS is unbiased when in-sample LD is available regardless of disease architecture (i.e. number of causals and distribution of effect sizes). We extend our method to use LD estimated from reference panels^16^ and show that a principal components based regularization of the LD matrix^17^ yields approximately unbiased and more consistent estimates of local heritability as compared to existing methods^6^.

We applied HESS to partition common SNP heritability at each locus in the genome using GWAS summary data for 30 traits spanning over 10 million SNPs and 2.4 million phenotype measurements. First, we show tht common SNPs explain a large fraction of the total familial heritability, ranging anywhere from 20% to 70% across the studied quantitative traits. Second, we showcase the utility of local heritability estimates in finding loci that explain more variance in trait than the top associated SNP at the locus (an effect likely due to multiple signals of association). Third, we contrast the polygenicity of all 30 traits by comparing the fraction of total SNP heritability attributable to loci with highest local heritability. We find that most of the 30 selected traits are highly polygenic with a small number of traits driven by a small number of loci. Finally, we report 55 “heritability hotspots” - regions of genome that have a high contribution to the heritability of multiple traits. Taken together, our results give insights into traits where further GWAS and/or fine-mapping studies are likely to recover a significant amount of the missing heritability.

## Materials and methods

### Overview of methods

We introduce estimators for the variance in trait explained by typed variants at a single locus (local heritability, 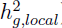) from summary GWAS data (i.e. Z-scores, effect sizes and their standard errors). We derive our estimator under the assumption that effect sizes at causal variants are fixed and genotypes are drawn from a distribution with a pre-specified covariance structure. The covariance, (i.e. pairwise correlation between any variants at a locus, LD) can be estimated in-sample, from the genotype data in GWAS, or from external reference panels (e.g. 1000 Genomes Project^16^). The finite sample size of the GWAS studies as well as the reference panels used to estimate LD induces statistical noise that needs to be accounted for to obtain an accurate estimation. Since the top projections make up the bulk of the summation, truncated-SVD lends itself as the appropriate regularization method to account for noise in the estimated LD matrices. Finally we extend our approach to consider multiple independent loci each contributing to the trait.

### Estimating heritability at a single locus from GWAS summary data

Let 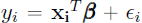, where *y_i_* is the trait value for individual *i*, X_i_ are the genotypes of individual *i* at the *p* SNPs in the locus, 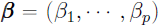) is the vector of fixed effect sizes for the *p* SNPs, and 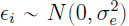 is the environmental effect. Assuming that β is fixed and X is random, the variance is

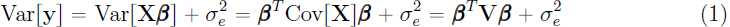

where V is a *p* × *p* variance-covariance matrix of the genotype vector (i.e. the LD matrix). If we make a standard assumption that the phenotypes are standardized (i.e. 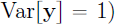), it follows that the amount of variance contributed by the p SNPs to the trait (i.e. local heritability) is 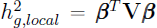. If the true effect size vector β and the LD matrix V are given, then computing 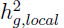 is trivial. In reality, however, the vector β is unknown and is estimated in GWAS involving *n* samples and *p* SNPs, where 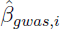 is estimated as the marginal standardized regression coefficient for SNP *i*

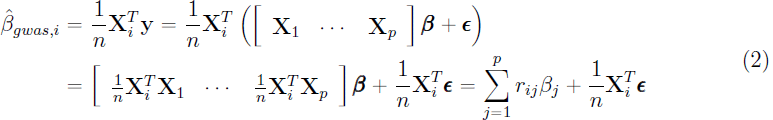

where 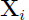 denotes standardized genotypes for SNP *i* across the *n* individuals, and 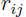 denotes the LD between SNPs *i* and *j*. Extending to *p* SNPs at the locus, if follows that 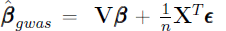 where V is the LD matrix. With β fixed and ∈ random, 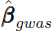 is a random variable with 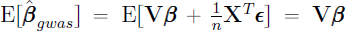, and 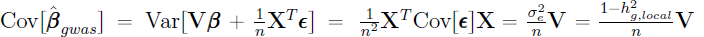. By central limit theorem, 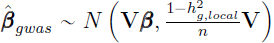.

As GWAS sample size (*n*) increases 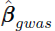 converges to 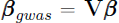. By simple substitution in Equation (1) it follows that an estimator for 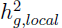 is

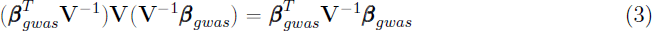

Unfortunately the finite sample size of GWAS induces statistical noise in the estimation of 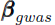 which leads to biased estimation if we would simply replace 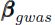 with 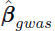 above, as 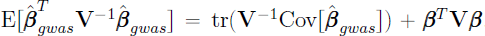. However we can correct for the bias 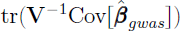 as follows.

Let 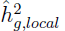 be an unbiased estimator of 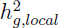, then by definition 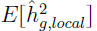 must satisfy 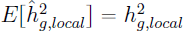. Then it follows that

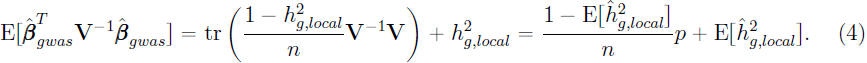

A sufficient condition for Equation (4) to hold is 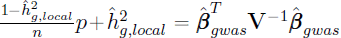. Solving for 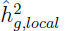 gives an unbiased estimator for 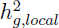

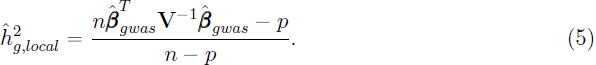

Following quadratic form theory^18^, the variance of 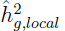 is

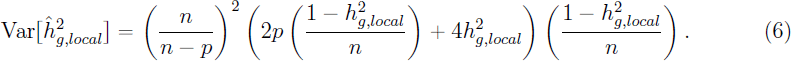

Since 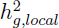, the true local heritability, is unknown, we use 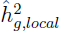 instead. For 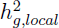 near 0, 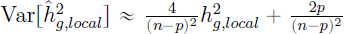 through Taylor expansion around 0. Thus, the plug in principle yields an estimation of 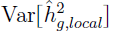 approximately equal to the truth in the expectation.

### Accounting for rank deficiencies in the LD

In the above derivation we made the assumption that the inverse of the LD matrix V exists. In practice, however, due to pairs of SNPs in perfect LD, V is usually rank deficient, and thus V^-1^ does not exist. In such cases we use the Moore-Penrose pseudoinverse **V**^†^. Let 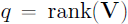, by rank decomposition, 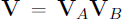, where 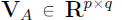 and 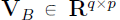 are matrices with full column rank and full row rank respectively, then 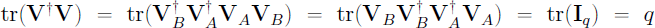. Accounting for rank-deficient LD matrix, we obtain an unbiased estimator, 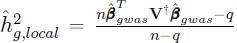. We make the same adjustment (replacing *p* with *q*) in the variance estimator for 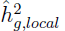.

### Adjusting for noise in external reference LD

When genotype data of GWAS samples is not available, we substitute the in-sample LD matrix V with external reference LD matrix 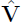 estimated from the 1000 Genomes Project^16^ using a population that matches the GWAS samples. However, due to limited sample size, external reference LD matrices contain statistical noise that biases our estimate. We apply truncated SVD regularization to remove noise from external reference LD matrix as follows.

First note that 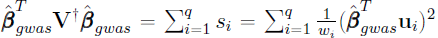, where 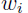 and u*_i_* are the eigenvalues and eigenvectors of the LD matrix V, and 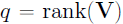. For external reference LD matrix 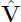 with eigenvalues and eigenvectors 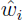 and 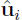, we have the same decomposition except that *s_i_* is replaced by 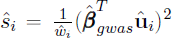. Through simulations, we show that most of the signal in 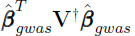 comes from *s_i_* where 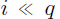 and that 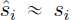 for 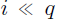 (see Supplementary Figure 1). In our previous works^19,20^, we propose to regularize 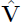 using ridge regression penalty. This regularization method is equivalent to replacing 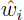 with 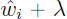, where λ is the penalty. We notice that a large λ is needed to drive down the noise (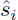 for large *i*), which diminishes the true signal at the same time. These results motivate us to apply truncated-SVD to remove noise in 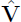, i.e. we estimate 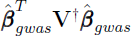 by 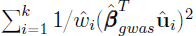, where 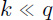. Let 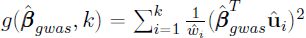, through eigen-decomposition of 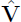, it can be shown that

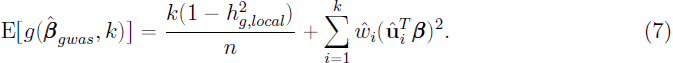

Since the true local heritability is 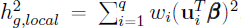, assuming 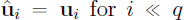, Equation (7) is an approximation of 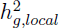 with bias 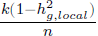. Correcting for this bias yields the estimator for the single-locus case

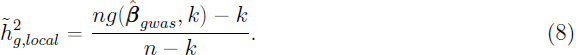

In theory, the variance of 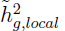 is 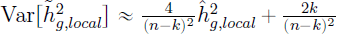. In practice, however, this gives an underestimation of the truth. Thus, we replace *k* with 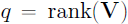.

### Extension to multiple independent loci

For genomes partitioned into m independent loci, the linear model for individual *i*’s trait value becomes 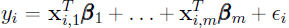 where 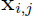 denotes the genotypes at the *p_i_* SNPs in the *i*-th locus for individual *i*, and 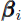 denotes the effect sizes of SNPs in this locus. Based on the revised model, we decompose Var[y] into

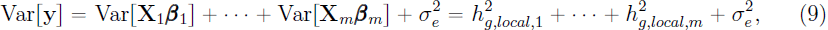

where 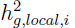 denotes the local heritability contributed by the *i*-th locus. In the case of multiple independent loci, the noise term 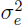 is equal to 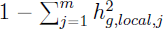. Thus, in order to correct for the bias generated by 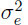, one need to know 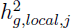 for all *j*. Accounting for bias and adjusting for noise in external reference LD 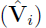 following strategies outlined in previous sections, we arrive at the estimator,

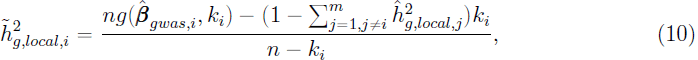

which defines a system of linear equations involving *m* variables 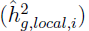 and *m* equations. A similar system of linear equations can be solved to obtain the variance estimator,

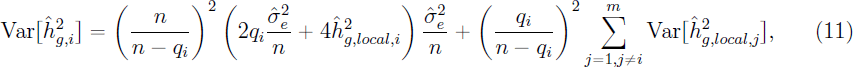

where 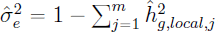

### Simulation framework

We use HAPGEN2^21^ to simulate genotypes for 50,000 individuals starting from half of the 505 European (EUR) individuals in the 1000 Genomes Project^16^ for SNPs with minor allele frequency (MAF) greater than 5% in one randomly selected region (1 Mb) on chromosome 1. We reserve the other half of the EUR individuals as external reference panel. From the simulated genotypes of the 50,000 individuals, we then simulate phenotypes based on the linear model 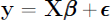, where **X** is the standardized genotype matrix with mean 0 and variance 1 at each column. To choose, β, the true effect size vector, we first select a subset *C* of 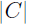 causal SNPs at random, such that 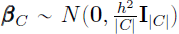, where *h*^2^ is the heritability to be simulated, and 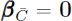. We draw ∈ from 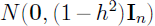 such that 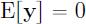, 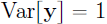, and that the SNP heritability for this locus is *h*^2^. For fixed β, we then generate replications of trait values **y** by varying ∈. Finally, we compute summary statistics, 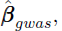 following procedures outlined in previous sections.

Empirical data sets

We obtained publicly available GWAS summary over European ancestry data for 30 traits from the websites of 11 GWAS consortia (see Table 1). For quality control, we restricted our analysis to GWAS studies involving at least 20,000 samples. We used the definition of independent loci as defined in^22^ (1.6 Mb on the average). To reduce statistical noise in LD matrix, we focused on estimating heritability attributable to common SNPs, i.e. SNPs with MAF greater than 5% in the European population. Prior to estimating heritability, we also removed SNPs with ambiguous alleles as compared to the reference panel (Supplementary Table 1) and applied our estimator as defined in Equation 11.

**Table 1:**
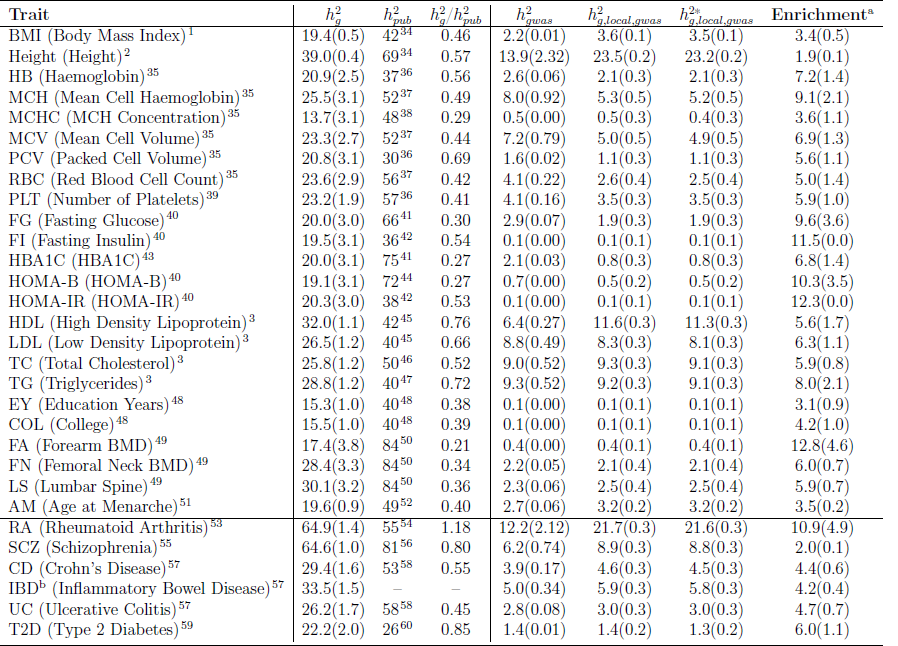
Total SNP heritability estimates and the amount of 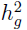 attributable to loci containing GWAS index SNPs 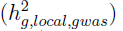 and index SNPs only 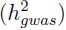. 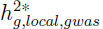 is the same as 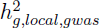 except that GWAS index SNPs are excluded in the computation. In supplementary, we report 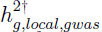, obtained by excluding all GWAS hits. We also report narrow sense heritability 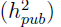 estimates obtained from twin or family studies. We list case-control traits where our estimate of 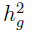 is biased due to ascertainment at the bottom of the table. ^a^Similar to^7^, we define enrichment as the ratio between the fraction of 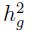 attributable to 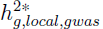 and the fraction of genome covered by these loci. We obtain standard errors by jackknife over the loci. ^b^IBD refers to the union of CD and UC.

Most GWAS apply the genomic control factor 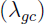 to the 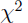 statistics to correct for inflation due to population structure^23^. However, recent works^6,24^ show that 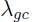 can not distinguish between inflation and true polygenicity and overestimates the correction factor needed for population stratification. Although dividing the 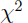 statistics by 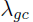 does not affect computing the ratios between local and genome-wide heritability^7^, it can result in underestimation of the total heritability across the entire genome (by scaling the local estimates of heritability in our framework). We notice all the summary GWAS data has at least one round of genomic correction. To account for this, we first re-estimate the 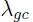 from summary GWAS and re-inflate the effect sizes with the estimated 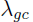 before estimating local heritabilities. To estimate 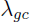 we conservatively regress 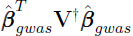 against 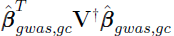 for the bottom 30% of loci with the smallest estimated local heritability. This is based on the observation that at loci where 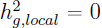, 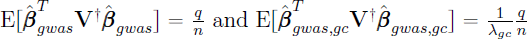, where 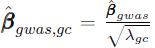 denotes GC corrected effect size vector. We report estimated 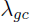 for all 30 traits in Supplementary Table 1.

We define GWAS hits as SNPs with p-values less than 5 × 10^-8^. For FI and HOMA-IR that do not have any GWAS hits at this threshold, we relax the threshold to 10^-7^. To avoid overestimation due to LD tagging, for each locus, we only select the most significant (i.e. smallest p-value) GWAS hit as the index SNP. Heritability attributable to index SNPs, 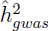, is then estimated as 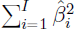, where 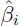 is effect size of the *i*-th index SNP, and *I* the number of index SNPs. We estimate the variance of 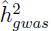 as 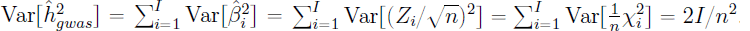.

For case-control traits, an adjustment factor is needed to correct for ascertainment^25^. We note that this adjustment factor is derived based on the infinitesimal model, and does not apply to our method, which assumes a fixed effect model. Therefore, we only report unadjusted heritability estimates for case-control traits. However, we note that ratio between local to genome-wide SNP heritability is not affected by this scaling factor.

## Results

### Performance of HESS in simulations

We used simulations to assess the performance of our proposed approach under a variety of disease architectures. First, we confirmed that by accounting for rank deficiency in the LD matrix we obtain unbiased estimation whereas the approach that uses the number of SNPs to correct for bias generated by the quadratic form^15^ leads to a severe under-estimation of heritability (Supplementary Figure 2). Second, we find that using the top 10-30 eigenvectors of the LD matrix (see Methods) provides a good approximation for the estimated heritability when LD is estimated from external reference panels (Supplementary Figure 1).

Next, we compared HESS to the recently proposed LD-score regression (LDSC)^6,7^ method that provides estimates of heritability from GWAS summary data. Although LDSC is not designed for local analyses due to model assumptions on polygenicity, it is able to estimate the variance in trait attributable to any sets of SNPs. As expected, in our simulations we find that LDSC is sensitive to the underlying polygenicity and, in general, yields biased estimation of heritability. In contrast, HESS provides an unbiased estimation of heritability across all simulated disease architectures when in-sample LD is available. For example, in simulations where 20% of the variants at the locus are causal explaining 0.05% heritability, HESS yields an estimate of 0.054% (s.e. 0.004%) as compared to 0.025% (s.e. 0.0009%) for LDSC (Figure 1). We attribute this to the fact that HESS does not make any assumption on the distribution of effect sizes at causal variants by treating them as fixed effects in the model. When LD from the sample is unavailable and has to be estimated from reference panels, both methods are biased with HESS yielding results closer to simulated heritability than LDSC (Figure 1 and Supplementary Figure 3).

**Figure 1:**
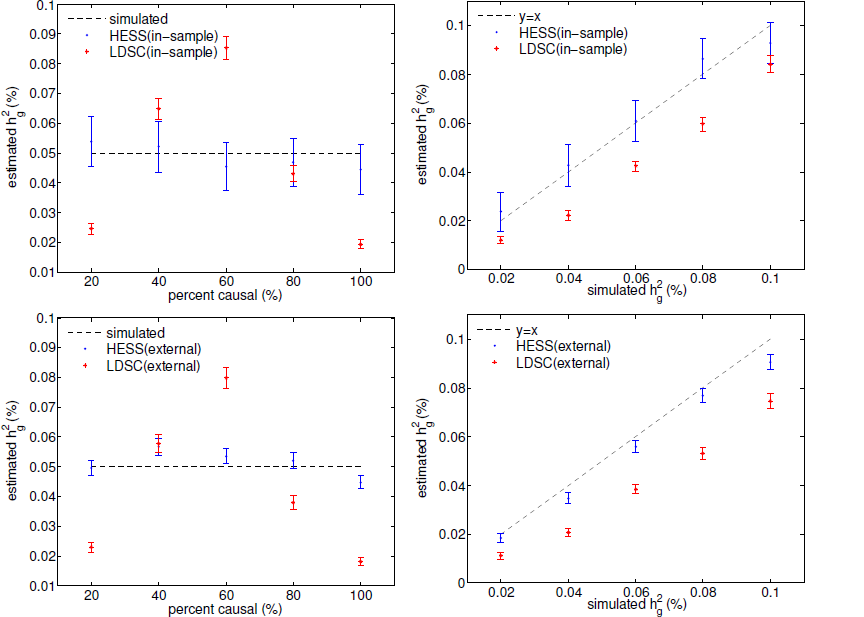
HESS provides superior accuracy over LDSC in estimating local heritability. HESS attains unbiased estimates when in-sample LD is used (top) and approximately unbiased estimates when reference LD is used (bottom). Mean and standard errors in these figures are computed based on 500 simulations, each involving 50,000 simulated GWAS data sets.

Finally, unlike LDSC that employs a jack-knife approach to estimate variance in the estimated heritability (thus requiring multiple loci), HESS provides a variance estimator following quadratic form theory (see Methods). We find that the variance estimator yields unbiased estimates when in-sample LD is available and under-estimates the theoretical variance when LD is estimated from reference data (Supplementary Figure 4). We attribute this to the fact that the rank of in-sample LD, which is computed based on many more samples, is in general larger than that of external reference LD.

### Common variants explain a large fraction of heritability

Having demonstrated the utility of HESS in simulations, we next applied our method to empirical GWAS summary data across 30 complex traits and diseases spanning more than two million phenotypic measurements (see Methods, Table 1, Supplementary Table 1). We estimated the local heritability at 1,703 approximately-independent loci^22^ using European individuals of the 1000 Genomes to estimate LD^16^. We first investigated the total contribution of common variants (MAF > 5%) to the heritability of complex traits. We summed up the local estimates provided by our method to obtain an estimate for the total genome-wide heritability for all genotyped SNPs. For traits where the SNP-heritability was previously reported we find a broad consistency between our estimate and the existing estimates from the literature (with differences likely due to different quality controls steps that retained slightly different SNP sets) (see Table 1). For example, HESS estimates a genome-wide SNP heritability 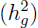 of 19.4% (s.e. 0.5%) for BMI and 39.0% (s.e. 0.4%) for height as compared to previously reported estimates of 21.6% (2.2%) for BMI^1^ and 49.8% (4.4%) for height^2^. Most importantly, we find that common SNPs explain a large fraction of the previously reported narrow sense heritability for all quantitative traits we interrogated ranging from 21% for Forearm BMD to 76% for HDL(Table 1). Although we observe a very high contribution of common SNPs to case-control traits as well, we note that our estimator can be biased due to ascertainment in this case (see Methods).

### Hidden heritability at known risk loci

Recent work^10,26^ has shown that the total heritability explained by all variants at the GWAS risk loci 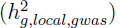 is higher than heritability explained by GWAS index SNPs 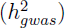. This suggests that a fraction of the missing heritability is due to multiple causal variants or poor tagging of hidden causal variants at known risk loci. We used HESS to quantify the gap between these two estimates of heritability at known risk loci. We find several traits for which 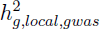 is signicantly larger than 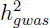. For example, 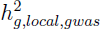 is almost two fold higher (24.2%, s.e. 0.2%) than 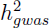 (14.3%, s.e. 0.01%) for height (Table 1). The difference can be accounted by incomplete tagging of hidden causal variant(s) or allelic heterogeneity (i.e., multiple causal variants). Indeed, conditional analysis identified 36 GWAS loci that contain multiple signals of associations (for a total of 87 GWAS risk SNPs at these loci) for height^27^. Restricting to the 43 loci that contain these multiple causal loci we estimate 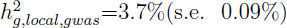, an almost 3-fold increase over 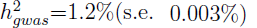. These loci, 8.7% of all GWAS loci for height, contribute to 38.1% of the difference between 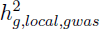 and 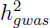 across all loci, thus suggesting that the difference is likely due to multiple signals of association. We observe similar results suggesting multiple signals of associations for HDL, TG, RA, SCZ (Table 1).

In contrast, the majority of traits show similar 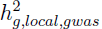 and 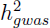 (see Table 1) suggesting a single causal variant at these loci very well tagged by the index GWAS variant. For example, it is known that LDL is strongly regulated by single non-coding functional variant at the SORT1 locus^3,28^ and that bone mineral density traits (FN) are strongly regulated by WNT16^29,30^. Finally, we note that reference panel LD approximation makes HESS is intrinsically conservative (see Methods) which can also lead to 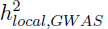 being less than 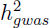.

Contrasting polygenicity across multiple complex traits

Most studied common traits exhibit a strong polygenic architecture (i.e. an abundance of loci of small effect contributing to trait)^1,2,3,9^. We recapitulate this observation using the HESS analysis and find a strong correlation between chromosome length and the fraction of heritability it explains for most traits we analyze here (Figures 2, 3). We also observe, consistent with previous findings^31^, regions such as FTO on chromosome 16 and HLA on chromosome 6 contributing disproportionately to the fraction of heritability for HDL, BMI, and RA, respectively.

**Figure 2:**
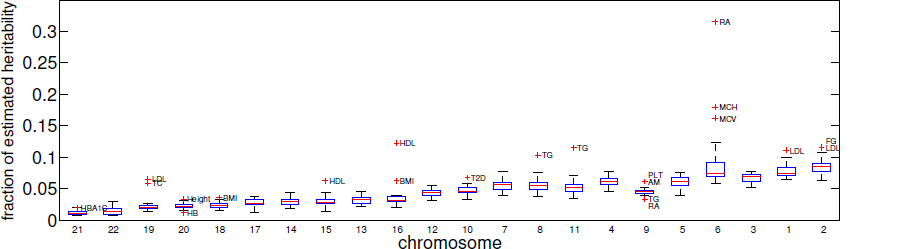
Fraction of 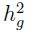 per chromosome across the 30 traits studied. Here, the chromosomal heritability is obtained by summing local heritability at loci within the chromosome. For each chromosome we plot the box plots of estimates at the 30 considered traits. Chromosomes are ordered by size. With some notable exceptions, all traits show a strong polygenic signature of genetic architecture.

**Figure 3:**
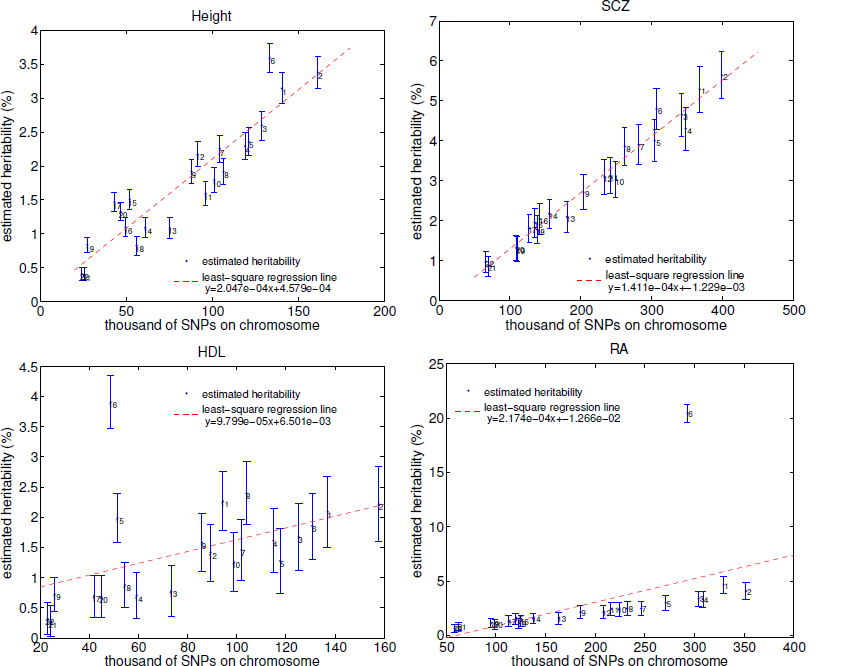
Heritability attributable to each chromosome for four example traits. The chromosomal heritability is obtained by summing local heritability at loci within the chromosome. Standard error is obtained by taking the square root of the sum of variance estimation. See Supplementary Note for results across all traits.

Next, we sought to quantify the variability in polygenicity across traits. We rank order loci based on their estimated local heritability, sum their contribution and plot it versus the percentage of genome they occupy (Figure 4). For highly polygenic traits, we expect the cumulative fraction of total SNP heritability to be proportional to the fraction of genome covered, whereas for less polygenic traits, we expect to see a small fraction of the genome accounting for a large fraction of total SNP heritability. For example, in schizophrenia and height the top 1% of the loci with the highest local heritability contribute to 3.9%(s.e. 0.94%) and 8.2%(s.e. 1.8%) of the total SNP heritability of these traits, respectively. This is consistent with previous reports on the degree of polygenicity of these traits^2,3,9^. At the other extremes, RA and lipid traits (HDL, LDL, TC, TG) have a lower degree of polygenicity, with the top 1% of loci accounting for 15-25% of the total SNP heritability. We note that the different degrees of polygenic signals across traits reflect both a difference in disease architecture (i.e. distribution of effect sizes) as well as a difference in the sample sizes for the GWAS summary data.

**Figure 4:**
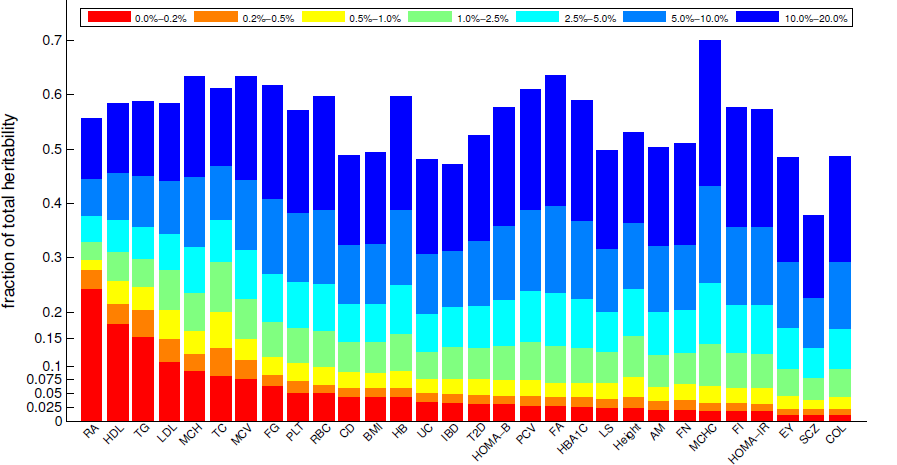
Stacked bar plot showing the percentage of total heritability attributable to different fractions of genome. We rank ordered all genomic loci by their explained heritability and quantified the fraction of total heritability attributable to different percentile ranges. Traits with high polygenicity tend to have bars with height proportional to bin size (e.g. Height and SCZ), whereas less polygenic traits tend to have bars much larger than bin size (e.g. RA and HDL).

A different perspective of polygenicity is to restrict to GWAS risk loci (as they clearly contain risk variants) and contrast the proportion of explained variance with the proportion of the genome they occupy. We observe a wide distribution across traits reflecting diverse genetic architectures as well as different sample sizes for the GWAS performed for these traits. For example, approximately 30% of loci across the genome harbor a risk variant for height and account for 50% to the total SNP heritability (an 1.6-fold enrichment). On the other hand, while only 5% of the loci contain GWAS risk variants for HDL, these loci collectively explain 25% of the SNP heritability of HDL (a 5-fold enrichment) (Figure 5).

**Figure 5:**
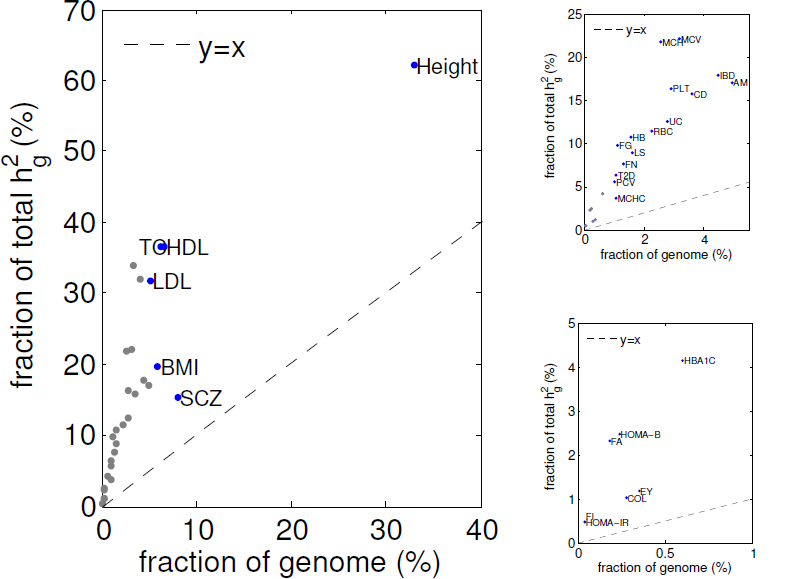
Fraction of 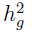 explained by all loci that contain a GWAS hit versus the fraction of genome covered by these loci. Images on the right focus successively on the traits near the bottom left.

### Loci that contribute to heritability of multiple traits

The 30 traits studied in this work share genetic correlations^6^ and we recapitulate this correlation using HESS estimates (Supplementary Figure 5). Motivated by this, we searched for pleiotropic loci which we define as loci that contribute more than 0.3% of the total SNP heritability for at least 3 out of the 30 analyzed traits. In total, we identified 55 such loci distributed genome-wide (see Figure 7 Supplementary Figure 6). As expected, the HLA region (chr16:26-34M), displays strong pleiotropic signal, particularly for immunologically relevant phenotypes (see Figure 7). For instance, locus chr16:32-33M contributes more than 0.3% of total SNP heritability for over 10 traits, with exceptionally strong signals for RA, UC, and IBD (see Figure 7). We also observe several pleiotropic loci on chromosome 10 and 16, including the loci chr10:63-66M, which contributes to PLT, IBD, CD, and UC, and the loci chr16:49-52M, which contributes to CD, IBD, and FN. Previous research has revealed the association between platelet count and inflammatory bowel diseases^32^ and the association between IBD traits and bone mineral density traits^33^. Our results suggest that the associations may be caused in part by the pleiotropic effect of the regions chr10:63-66M and chr16:49-52M, respectively. We note that the selection of traits can bias the identification of pleitropic loci towards over-represented traits. Nevertheless, local heritability analysis is still a useful tool to quantify the fraction of total SNP heritability contributed by a single loci and provide valuable insights into identifying pleiotropic loci.

**Figure 6:**
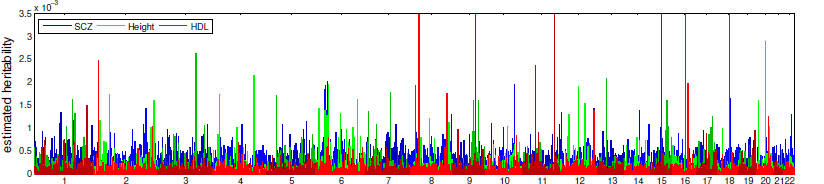
Manhattan-style plots of regional heritability across the genome for the trait Height, HDL, and SCZ. See Supplementary Note for results across all traits.

**Figure 7:**
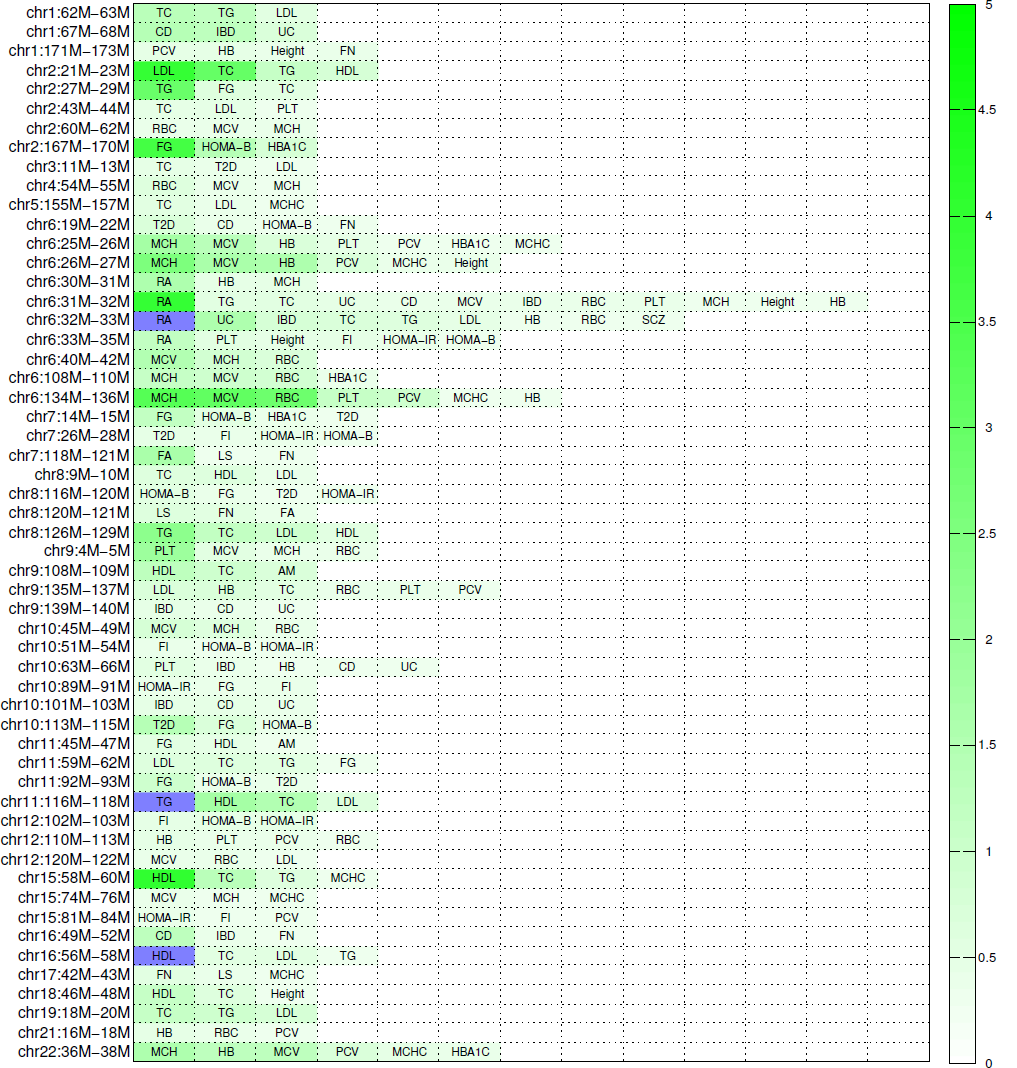
Heat map showing the fraction of total SNP heritability (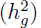) contributed by each of the 55 “pleiotropic” loci. For each locus, we only display the traits to which the locus contributes more than 0.3% of total SNP heritability. We mark traits to which the locus contributes more than 5% of the total SNP heritability in dark blue.

### Discussion

We have presented HESS, an unbiased estimator of local heritability from GWAS summary data. Our approach extends existing works that treat SNP effects as fixed and model genotypes as random variables^15^ - we proposed a method to regularize LD matrix estimated from external reference panel and generalized the estimator to multiple independent loci. Through extensive simulations, we demonstrated that HESS yields more consistent and less biased local heritability estimates than another recently proposed method that models SNP effects as random variables. We applied HESS on GWAS summary data of 30 traits from 11 GWAS consortia and showed that our results recapitulate previous findings. We then used the local estimate to contrast polygenicity of these traits, find loci with multiple causal variants, and identify heritability hot spots. We note that the enrichment of heritability at GWAS risk loci could be leveraged into prioritizing GWAS or fine-mapping; for example, traits with small enrichment of heritability at GWAS risk loci are more suitable for larger GWAS, whereas traits with large enrichment of heritability at known risk loci could be investigated further through fine-mapping.

Our focus in this work is estimating local heritability attributable to common SNPs (MAF > 5%). Thus, we did not explore the issues of effect size and LD at rare variants. We also note that methods that adjust heritability estimation for case-control traits are currently lacking. Our reported heritability estimation for case-control traits can be biased due to ascertainment in GWAS for case-control traits. Nevertheless, our method still demonstrates its utility in studying and comparing genetic architectures of complex traits. We conjecture that future work that addresses local heritability estimation including both common and rare variants as well as adjustment of local heritability under the fixed-effect model will further improve the utility of our approach.

## Acknowledgements

Acknowledgement

We are very grateful to Alkes Price, Yakir Reshef, Brielin Brown, and Nicholas Mancuso for their helpful discussions and feedback that greatly improved the quality of this manuscript. The authors declare no conflict of interest.

